# Spatial learning in multi-scale environments: Roles of hippocampus, orbitofrontal cortex, and retrosplenial cortex

**DOI:** 10.1101/2025.10.20.683540

**Authors:** Yidan Qiu, Senning Zheng, Huakang Li, Shuting Lin, Ruiwang Huang

## Abstract

Navigation cognition involves the learning of multi-scale environments and the formation of cognitive maps. How do humans build cognitive maps in multi-scale environments? Cognitive maps are thought to be organized hierarchically, with local representations for subareas and global representations for the entire environment. However, it remains unclear how spatial learning influences the representations of multi-scale environments and their underlying neural mechanisms across different levels of representation. In the current study, we built a virtual environment (VE) with four orthogonally positioned rectangular rooms, each containing eight objects at the corners. Twenty-three healthy subjects completed a four-session spatial memory experiment conducted over two weeks. We measured their brain activity by using BOLD-fMRI at two stages: pre-learning stage and post-learning, when they were judging the relative direction between the objects within the VE. We found that with the progression of learning, the subjects shifted from relying on local, directional cues to using more global representations of the environment. At the neural level, the hippocampus (HIP), retrosplenial cortex (RSC), and orbitofrontal cortex (OFC) played distinct roles in encoding spatial information across the two learning stages. Specifically, after learning, the HIP shifted from local to global representations, while the RSC and OFC supported the integration of spatial information across these representational levels. In addition, the anterior cingulate cortex was involved in forming global representations, facilitating efficient spatial processing as learning advanced. These findings revealed how spatial learning leads to adaptive shifts in brain activity, contributing to the formation of cognitive maps in complex, multi-scale environments.

**Highlights:**

- Using fMRI to study neural mechanisms of cognitive map formation in a multi-scale environment.
- Detected that local and global spatial representations coexist and evolve during spatial learning.
- Found that learning progress is associated with activation changes in the entorhinal cortex, hippocampus, and retrosplenial cortex.
- Revealed the complementary roles of hippocampus and orbitofrontal cortex in representing the environment.
- Showed that the retrosplenial cortex improves both local and global representations, supporting the integration of multi-scale spatial information.

## 1. Introduction

Imagine navigating a large amusement park with various themed areas, each containing different rides and attractions. To get around, you need to understand both the layout within each area (i.e., local reference frame, RF) and the connections between the areas across the entire park (i.e., global RF). This navigational ability relies on cognitive maps, which are mental representations of spatial environments that help us to know where we are, to plan routes, and to make navigation decisions ^1,2^. Cognitive maps are essential for navigating complex environments with multiple subareas and landmarks, as they integrate information across RFs ^3,4^. Beyond navigation, cognitive maps support broader cognitive functions such as episodic memory and decision-making ^5,6^, enabling efficient inferences from limited experience, guiding novel decisions, and supporting flexible behavior ^4^.

How are cognitive maps organized in multi-scale environments? This issue has been studied using behavioral ^7–9^ and neuroimaging ^10–12^ experiments. A widely accepted framework is that cognitive maps are structured hierarchically in multi-scale environments, with three levels of representation: local, categorical, and global ^13–15^. Local representations capture spatial relationships within smaller areas (e.g., individual rooms), enabling navigation within specific areas ^13,16^. Categorical representations can distinguish boundaries and characteristics between areas, supporting transitions across different areas ^17^. Global representations provide a unified view of the entire environment, using global coordinates to represent spatial relationships across all areas, thus facilitating flexible navigation ^17–19^. The organization of cognitive maps is influenced by learning ^16,20^. Some studies ^21,22^ suggested that individuals initially rely on local representations, gradually incorporating local spatial information into a global framework. Conversely, other studies ^23,24^ proposed that global representations are build first, and are then refined into hierarchical structures to capture detailed local relationships. Based on the hierarchical organization framework, the current study aims to reveal how spatial learning influences the development and integration of local and global representations.

How does the brain support the formation of cognitive maps in multi-scale environments? Previous studies ^1,25,26^ identified several brain regions involving in spatial representation, such as the hippocampus (HIP), entorhinal cortex (EC), orbitofrontal cortex (OFC), and retrosplenial cortex (RSC). Specifically, the HIP is essential for encoding memory, spatial distances, and location information, particularly through place cells ^24,27,28^. The EC is critical for providing a coordinate system through grid cells, which helps in the formation of cognitive maps ^2,6^. Together, the HIP and EC support the spatial coding necessary for navigation and cognitive mapping ^25,29^. The OFC is believed involving in cognitive mapping to support flexible decision-making, particularly when tasks require integrating local and global information ^30,31^. The OFC is also engaged in planning routes during navigation ^29^. The RSC is involved in anchoring cognitive maps to the environment and integrating different perspectives for a location ^25,32^. Moreover, the RSC has been found to represent location and facing direction in multi-scale environments ^33^, suggesting its role in linking local and global RFs. Collectively, these regions are involved in building cognitive maps of novel environments ^34,35^.

The present study aimed to understand how cognitive maps are built and updated during spatial learning in multi-scale environments. Following the paradigm of Marchette et al. ^33^, we invited young adult healthy subjects to perform a judgement of direction (JRD) task in a virtual environment containing subareas (Fig. 1), requiring the subjects to integrate local, categorical, and global RFs. We hypothesized that as learning progresses, the subjects would shift from relying on local to more global spatial representations, with corresponding changes in brain activity across key regions such as the RSC, HIP, EC, and OFC. To track the updating of cognitive maps, we extended this approach with a multi-stage learning design, incorporating two task-fMRI sessions: a pre-learning stage and a post-learning stage (after extensive learning). This design allowed us to examine the changes in both behavior and brain activity as the subjects became familiar with the environment. In the behavioral analysis, we examined the influence of different levels of spatial information on task performance across the two learning stages, revealing how reliance on local, categorical, and global representations evolved. In the fMRI analysis, we identified the roles of key brain regions, including the RSC, HIP, EC, and OFC, in supporting local and global representations during learning. Our study provides new insights into the neural mechanisms underlying cognitive maps and the interaction between hierarchical and map-like representations in complex environments.

**Fig. 1.**
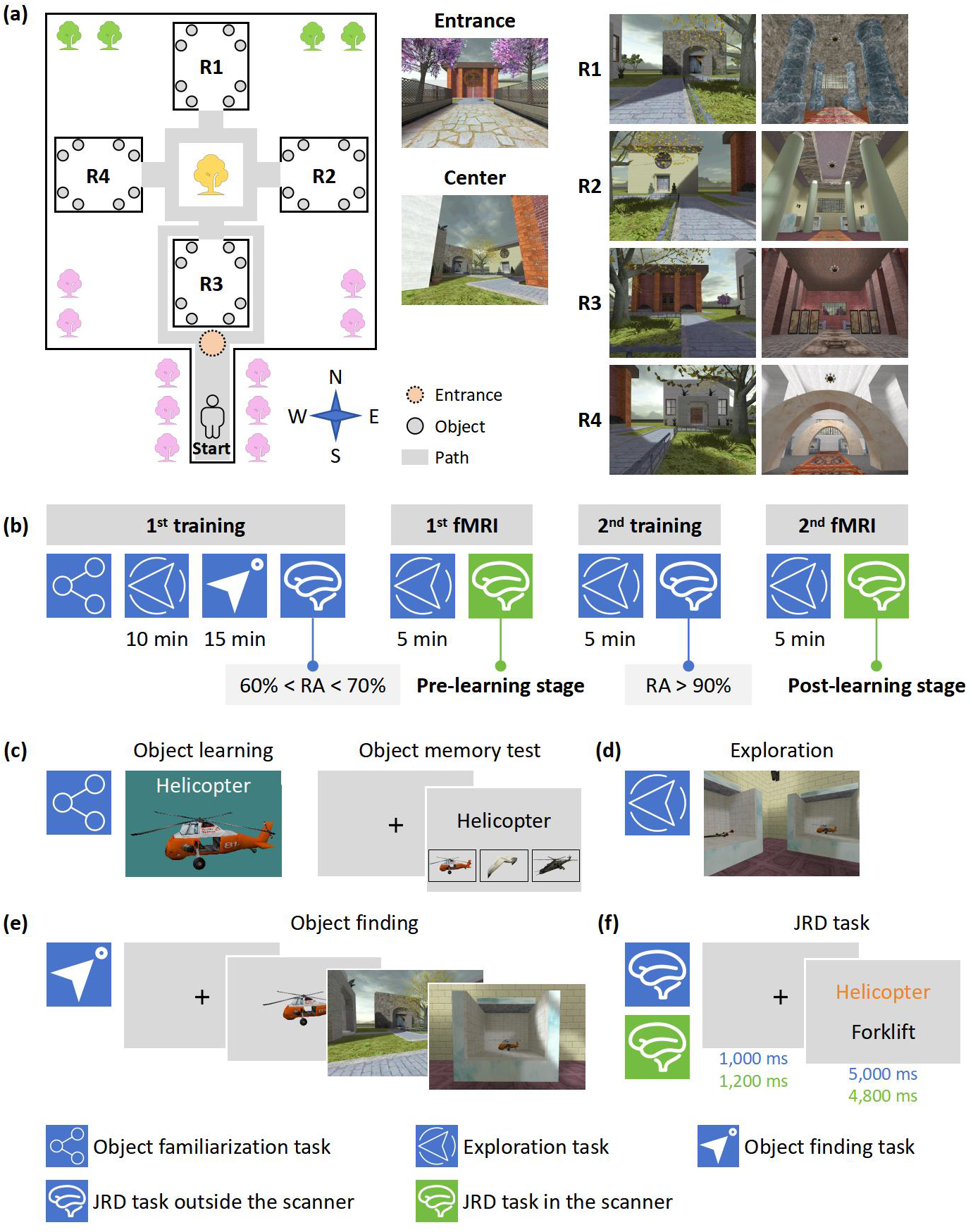
Overview of the experimental tasks. **(a)** Virtual environment (VE). (Left) An aerial view of the park with four rooms, R1-R4, arranged orthogonally. (Middle) Subject’s perspective at the entrance and center of the VE. (Right) Subject’s perspective from outside and inside the rooms. The directions N, E, S, and W represent north, east, south, and west, respectively. **(b)** Experimental timeline. The experiment was conducted over 4 non-consecutive days. The intervals between each behavioral training and fMRI sessions was ≤ 2 days, while the interval between the 1^st^ fMRI scan session and the 2^nd^ behavioral training session was ≤ 8 days. RA = response accuracy. **(c)** Object familiarization task. Subjects learned about the objects used in the VE through an object learning phase and an object memory test. **(d)** Exploration task. Subjects freely explored the VE to learn its layout and object locations. **(e)** Object finding task. Subjects were instructed to locate specific objects within the VE. **(f)** Judgment of relative direction (JRD) task. Subjects judged the direction of one object relative to another. The blue icons indicate the tasks being conducted outside the scanner, and the green icon indicates a task being conducted inside the scanner.

## 2. Methods

### 2.1 Subjects

Twenty-five healthy adults initially enrolled in a 4-day spatial memory experiment (Fig. 1). One subject withdrew from the study, and another did not meet the required accuracy threshold in the second behavioral training session, resulting in a final sample of 23 subjects (11 F/12 M; mean age = 21.61 years old, range = 18-25 years old). All of the subjects had normal or corrected-to-normal vision and no history of neurological or psychiatric disorders. The study was approved by the Institutional Review Board (IRB) of South China Normal University (SCNU). Written informed consent was obtained from all subjects prior to the experiment, and they were compensated for their participation.

### 2.2 Stimuli

A virtual environment (VE) was created using the Source SDK Hammer Editor implemented in *Counter-Strike* (https://www.counter-strike.net/workshop/workshop). Fig. 1a shows that the VE consisted of a park with four rectangular rooms positioned in the north, east, south, and west. Each room contained eight objects (two objects in each corner), for a total of 32 objects in the four rooms. The objects were of four types, animals, plants, vehicles, and furniture, each type including eight objects. The placement of objects was randomized, and the objects of different types were evenly distributed across the four rooms to minimize memory effects based on the object types. No two objects of the same type were placed in similar locations across different rooms. According to the spatial scales, we defined (1) a global RF for the entire park based on the cardinal directions, and (2) a local RF for each room based on the positions relative to specific landmarks (door, opposite wall, left, or right) (Fig. 2). Thereby, the VE involves three hierarchical levels (Fig. 2): (1) local level, which represents the spatial relationships between objects within a room based on the local RF, (2) categorical level, which represents the rooms as separate spatial units, and (3) global level, which represents the spatial layout of the entire park using the global RF.

**Fig. 2.**
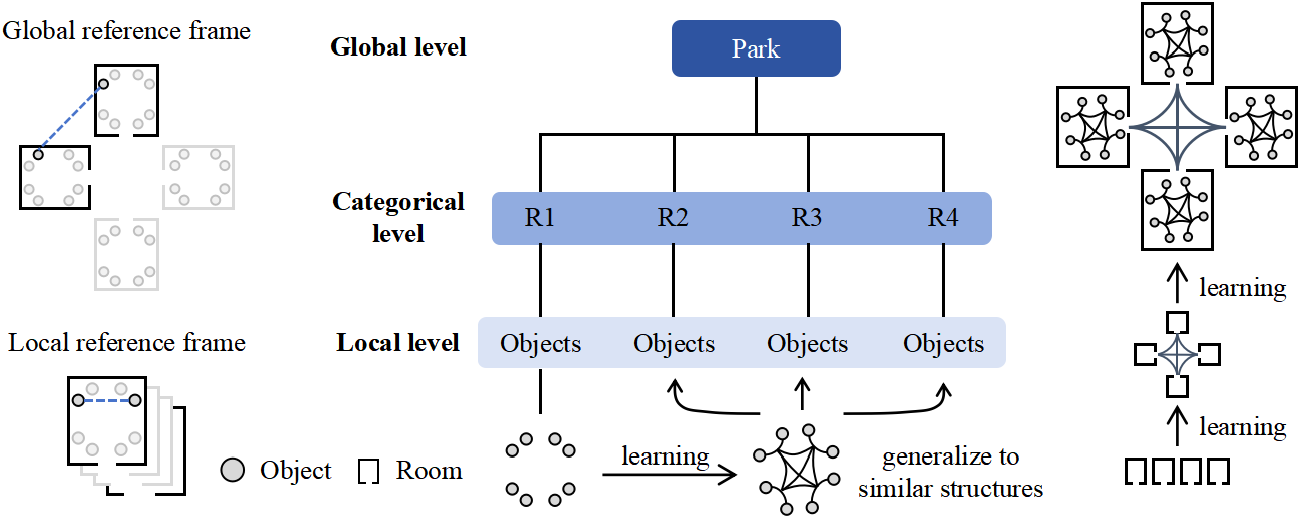
Illustration of the relationships among the three hierarchical levels, reference frames (RFs), and the learning process in the experiment. The local level of the environment captures spatial relationships between the objects within each room, based on the local RF. After learning, the subjects may strengthen and generalize these relationships to similar spatial structures. The categorical level reflects the semantic distinction between rooms (R1-R4). The global level represents the spatial layout of the entire park using the global RF, with spatial relationships across rooms becoming clearer after learning.

### 2.3 Experimental tasks

Experimental tasks were programmed in E-prime 2.0 (Psychology Software Tools, Pittsburgh, PA, USA).

#### 2.3.1 Object familiarization task

The subjects familiarized themselves with the 32 objects by reviewing a document that contained the names and pictures of each object (Fig. 1c). Once they felt prepared, they were requested to complete a memory test about the objects on a desktop. In each trial of the memory test, the subjects were shown the name of an object and were asked to select the corresponding picture from three options (Fig. 1c). They were required to achieve accuracy ≥ 95% (i.e., no more than a mistake).

#### 2.3.2 Exploration task

The subjects moved freely through the rooms and observed the 32 objects to learn their locations in the VE (Fig. 1d). The subjects controlled the first-place perspective using the mouse and adjusted the movement direction by pressing the arrow keys on the keyboard. They were asked to enter each room at least once. The task lasted approximately 10 minutes.

#### 2.3.3 Object finding task

The object finding task was conducted on two desktops, one for displaying the target objects (referred to as the “presentation desktop”) and the other for displaying the VE navigation (referred to as the “navigation desktop”). This setup allowed the subjects to view the indicator of target objects and to navigate within the VE simultaneously, without the indicator affecting their perspective.

The task involved the learning of 32 objects, each being presented twice in a pseudo-random order, resulting in a total of 64 trials. In each trial, a picture was shown on the presentation desktop to indicate the object to be located (Fig. 1e). On the navigation desktop, the subjects used the mouse and keyboard arrow keys to navigate and position themselves in front of the object. Once the subjects reached the object, they pressed the space bar on the presentation desktop to trigger the appearance of the next target object. There was no time limit for the subjects’ responses, allowing them ample time to locate the target object.

These trials were organized into 8 blocks, each block containing 8 trials. In each block, two objects from the same room were presented consecutively, and the order of room presentation was randomized. For example, using A-D to represent the four rooms and 1-8 to represent the 8 objects in each room, a possible trial order in a block could be “A1-A2-C4-C7-D6-D4-B2-B8”. The entire task lasted about 15 minutes. The object finding task was designed to reinforce the subjects’ memory of the object locations, facilitating the acquisition of both the local spatial relationships between the objects within each room and the global relationships between the rooms in the VE.

#### 2.3.4 Judgement of direction (JRD) task

In the JRD task, the subjects were required to judge the relative directional relationship between two objects within the same room. The task consisted of 64 trials and was conducted only on the presentation desktop. Each trial began with a 1 s fixation cross, followed by the presentation of two object names on the screen. The subjects were asked to imagine themselves standing in front of the first object (the reference object) and to judge whether the second object (the target object) was located to their left or right. They had 4.8 s to respond by pressing either the key of “1” (for left) or “2” (for right) on the keyboard. Both the objects were located in the same room. Additional 16 catch trials were randomly inserted to the JRD task to test whether the subjects were actively retrieving the object locations in the park. In these catch trials, only an object name was shown on the screen and the subjects were asked to identify which room contained the object. They had 4.8 s to respond by pressing either the “1”, “2”, “3”, or “4” key for the four rooms on the keyboard. No feedback was provided during the task.

### 2.4 Experimental procedures

Fig. 1b shows the experimental procedures that were conducted over four sessions in 2 consecutive weeks: (1) the 1^st^ behavioral training session, (2) the 1^st^ fMRI scan session (pre-learning stage), (3) the 2^nd^ behavioral training session, and (4) the 2^nd^ fMRI scan session (post-learning stage). The interval between each training and scan sessions was less than 2 days and the interval between the two training sessions was about a week.

In the 1^st^ behavioral training session, the subjects first performed the object familiarization task (Fig. 1c), and then entered the VE for object location learning by completing the exploration task (Fig.1d) and the object finding task (Fig. 1e). Next, the subjects performed the JRD task. If their accuracy was < 60%, they repeated the object finding task before performing the JRD task again, until they achieved accuracy ≥ 60% in the JRD task. To prevent ceiling effects, we excluded any subject who achieved accuracy ≥ 70% in JRD task of the 1^st^ behavioral training session from further experiments. No subject exceeded this threshold.

The 1^st^ fMRI scan session included six runs of the JRD task (Fig. 1f). Each run consisted of 64 JRD trials and 4 catch trials. Before the fMRI scan, the subjects completed a 5-minute exploration task outside the scanner to consolidate memory of the object locations. Following the method of Marchette et al. ^33^, we used only the objects from two orthogonally-aligned rooms in the 1^st^ fMRI scan session. The objects from the remaining two rooms were reserved for the 2^nd^ fMRI scan session. In each fMRI run, the duration of the fixation cross was adjusted to 1.2 s to match the fMRI acquisition timing. The object names were displayed for 4.8 s, which is the response window for both the JRD and catch trials. If the subjects responded within the response window, the trial screen remained on display until the end of the response window. The next trial began immediately after the response window. After the subjects completed each fMRI run, we requested them to report no sleep during the scanning. After the subjects completed the six runs in about 40 min, they took a 10 min break outside of the scanner.

In the 2^nd^ behavioral training session, the subjects first performed the object finding task, followed by the JRD task. This process was repeated until their accuracy in the JRD task was increased to at least 90%. In the 2^nd^ fMRI scan session, the subjects followed the same procedure as the 1^st^ fMRI scan session but the objects come from the remaining two rooms. The subjects first completed a 5-minute exploration task before the fMRI scan, then completed 6 runs of the JRD task during the scan.

### 2.5 MRI data acquisition

All imaging data were collected on a Siemens Trio 3T MRI scanner equipped with a 32-channel phased-array head/neck coil. The functional MRI (fMRI) data were acquired using a single-shot, simultaneous multi-slice (SMS) gradient-echo echo-planar imaging (EPI) sequence with the following parameters: repetition time (TR) = 1,200 ms, echo time (TE) = 41.6 ms, flip angle = 52°, slice acceleration factor = 5, field of view (FOV) = 211 × 211 mm^2^, data matrix = 88 × 88, slice thickness = 2.4 mm without inter-slice gap, voxel size = (2.4 mm)^3^, anterior-to-posterior phase encoding direction, and 65 transversal slices covering the whole brain. To correct for susceptibility-induced geometric distortions and BOLD signal loss, we also acquired a field map by using a double-echo gradient-echo sequence with TR = 735 ms, TE1/TE2 = 5.04 ms/7.50 ms, flip angle = 60°, FOV = 211 × 211 mm^2^, voxel size = (2.4 mm)^3^, and 65 axial slices. High-resolution anatomical images were obtained using a T1-weighted 3D MP-RAGE sequence with TR = 1,600 ms, TE = 2.98 ms, flip angle = 9°, slice thickness = 1 mm, FOV = 256 × 256 mm^2^, data matrix = 256 × 256, voxel size = (1.0 mm)^3^, and 176 sagittal slices covering the whole brain. For each subject, all the imaging data were obtained in the same session.

During the fMRI scan, the subjects viewed the stimuli on a screen via a mirror mounted on the head coil. Behavioral responses were recorded using a 4-button bimanual response box. Foam padding was used to minimize head movement, and earplugs were provided to reduce acoustic noise. To enhance BOLD signal quality in the OFC and temporal cortex, each subject’s head was tilted backward at an angle of 15-30° by placing a towel under the neck during scanning ^36^. The MRI data were preprocessed using fMRIPrep 20.2.7 ^37^. The detailed pre-processing procedure is described in Supplementary Materials.

### 2.6 Definition of the regions of interest (ROIs)

We selected eight ROIs, including the bilateral HIP, EC, OFC, and RSC. The HIP was defined according to the Harvard-Oxford subcortical structural atlas ^38^ with a probability ≥ 50%. The EC was defined according to the Juelich Histological Atlas ^39^, also with a probability ≥ 50%. The OFC was defined according to the Brainnetome Atlas ^40^, which provides a fine-grained parcellation based on functional and structural connectivity. The RSC was defined as Brodmann’s Areas (BA) 29 and 30 within the posterior cingulate cortex (PCC) ^41^. We also selected the bilateral primary motor cortex (M1) as the control ROIs. The M1 was defined according to the Juelich Histological Atlas ^39^ with a probability ≥ 50%.

### 2.7 Linear mixed-effects model (LMM) for behavior analysis

Linear mixed-effects models were used to assess how spatial information influenced subjects’ response time (RT) across two different learning stages. Specifically, we built two candidate models, LMM1 and LMM2, by using the same fixed-effect terms. Their difference was that LMM1 (Eq. 1) did not include the learning stage as a random effect, while LMM2 (Eq. 2) included it as a random effect.

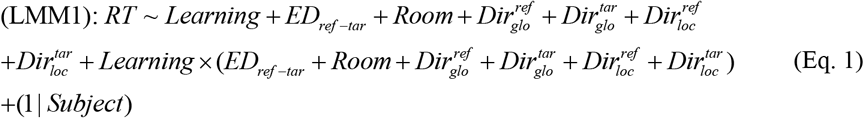

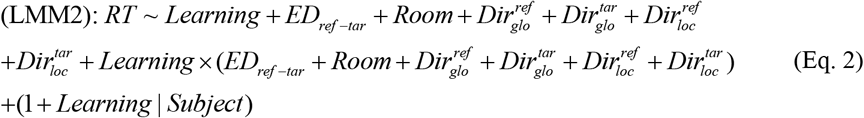

where *Learning* is a categorical variable denoting the two learning stages (i.e., the pre-learning and post-learning stages). *ED*_*ref-tar*_ is Euclidean distance between the reference (*ref*) and the target (*tar*) objects (Fig. S1 in Supplementary Materials). Because the reference object and the target object were in the same room in all of the trials, the *ED*_*ref-tar*_ is the same under the local and global RFs. 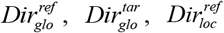 and 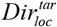 are categorical variables representing the 4 possible directions (north, east, south, and west), denoting the global (*glo*) or local (*loc*) facing directions of the reference (*ref*) or target (*tar*) objects (Fig. S1 in Supplementary Materials). Because only two orthogonally-aligned rooms were included in each of the fMRI sessions, we set the *Room* variable as a binary factor, indicating whether the room was aligned along the north-south or east-west axis. *Learning* × (·) denotes the interaction between the learning stage and other variables, (1| *Subject*) in Eq.1 denotes the random intercept for each subject, and (1+ *Learning* | *Subject*) in Eq.2 represents the random intercept and the random slope for the learning stage within each subject.

All the LMMs were conducted using the lme4 package in *R* ^42^. The values of RT were log-transformed prior to modeling to meet the assumption of normality. The models were compared using the Akaike Information Criterion (*AIC*), with lower *AIC* values indicating better model fit.

### 2.8 General linear model (GLM) for fMRI data

We conducted both the ROI-based and whole-brain voxel-wise GLM analyses to identify brain regions involving in the JRD task. The GLM analyses were carried out with FSL (version 6.0.5.1). At the subject-level, the GLM included three regressors: (1) the JRD trials, (2) a parametric modulator for the *ED*_*ref-tar*_, and (3) catch trials. In addition, six nuisance regressors for head motion parameters (three translations and three rotations) were included to control for motion-related artifacts. For each subject, we estimated the *β*-values associated with the JRD trials and the *ED*_*ref-tar*_.

At the group-level, we performed a random-effects analysis to assess the effects of the JRD trials and the *ED*_*ref-tar*_ on brain activation across both learning sessions. Gender and age were included as covariates to control for potential demographic effects. Specifically, we conducted an ROI-based analysis to evaluate activation patterns related to the JRD trials and the influence of *ED*_*ref-tar*_ within the pre-defined brain regions. A paired-sample *t*-tests was used to examine the differences in brain activation between the pre- and post-learning stages for the JRD trials and the *ED*_*ref-tar*_ separately. In addition, we also performed the voxel-wise GLM analysis in the whole-brain to identify other brain regions with differential activation between the two stages.

A brain-behavior correlation analysis was performed to examine the relationship between brain activation during the JRD task and individual differences in the learning process. For brain functional data, we first created spherical masks with a radius of 4 mm centered at the peak coordinates of significant clusters identified in the ROI-based GLM analysis. From these masks, we extracted the average *β*-values associated with the JRD trials and the *ED*_*ref-tar*_ from the subject-level GLM. For behavioral data, we extracted the random slopes for the leaning stage (denoted as *η*-values) from the LMM with lower *AIC* value for each subject. These *η*-values represent the effect of the learning stage on RT for each individual, capturing variability in learning effects across the subjects. Finally, we calculated Pearson’s correlation between the *η*-values and the extracted *β*-values.

### 2.9 Representational similarity analysis (RSA) for fMRI data

We performed RSA to determine whether brain regions encoded the local and global RFs at different learning stages (Fig. 2). The RSA was conducted using python packages, including NiBabel and SciPy. For each subject, two theoretical representational dissimilarity matrices (RDMs) were built to capture the spatial relationships of the reference objects at the local and global RFs during each learning stage (Fig. 4a). Because each room contained 8 objects and each learning stage used two rooms (containing 16 objects in total), the theoretical RDMs were 16 × 16 matrices. To build the local RDM, we aligned the four rooms within the VE according to their local spatial structures, i.e., rotating the rooms to unify the orientation of the doors and stacking them (Fig. 4a). Then, we computed the Euclidean distance between the 16 reference objects under the local RF 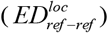 to capture their spatial relationships in a localized context, regardless of the room’s global orientation. In contrast, the global RDM was built to capture spatial relationships of the objects across the entire VE by using the Euclidean distance between the global coordinates of the 16 reference objects 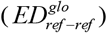 (Fig. 4a). We also built a categorical RDM to capture whether the objects were located in the same room or not (Fig. S2 in Supplementary Materials). The objects in the same room were defined as similar, and the objects in different rooms were considered dissimilar. However, due to the high inherent similarity between the global and categorical RDMs, we chose to exclude the categorical RDM from subsequent RSA.

**Fig. 3.**
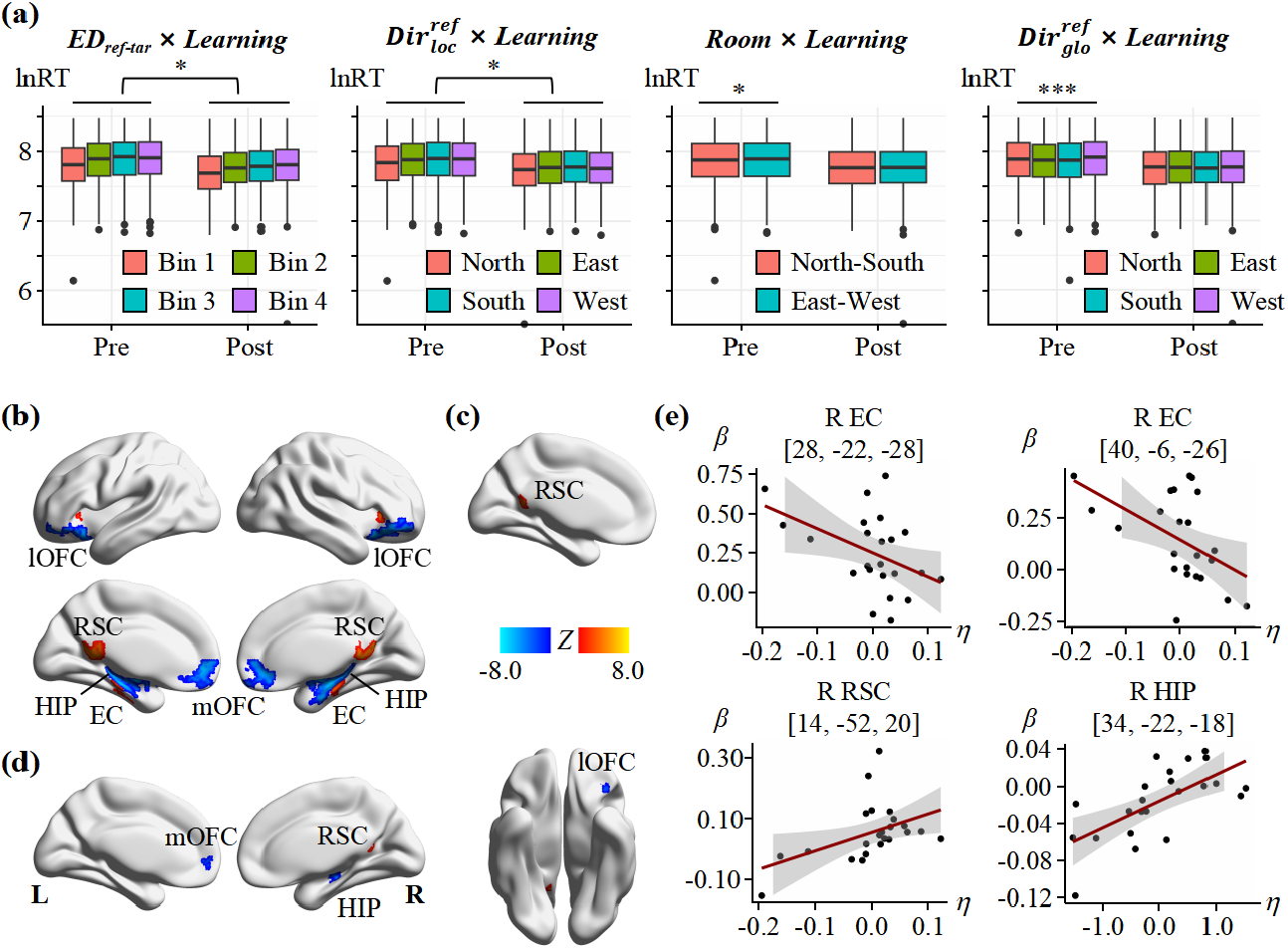
Behavioral performance and brain activation in the judgement of relative direction (JRD) task. **(a)** Interaction between spatial information and learning stage on response time (RT). *ED*_*ref-tar*_ represents the Euclidean distance between the reference and target objects, which was divided into four equal bins based on the quartiles of the distance distribution for ease of visualization. Bin1, Bin2, Bin3, and Bin4 represent increasing distances from closest to furthest. 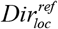 represent the local directions of a reference object. Because only two orthogonally-aligned rooms were included in each fMRI session, we set the *Room* variable as a binary factor, indicating whether the room was aligned along the north-south or east-west axis. 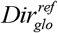 represent the global direction of a reference object. The boxplots indicate the interquartile range (IQR), with horizontal lines within the boxes representing the median, whiskers extending to 1.5 times the IQR, and dots showing outliers beyond 1.5 times the IQR. Asterisks above each learning session denote the main effect of spatial information, and asterisks across learning stages indicate interaction effects between learning stage and spatial information. *, *p* < .05, **, *p* < .01, ***, *p* < .001. The results of post-hoc analysis are provided in Table S1 (Supplementary Data). **(b)** Brain regions with significant activations in the JRD task. **(c)** Significant activation in the left retrosplenial cortex (RSC) for the contrast of post-learning > pre-learning stage in the JRD trials. No regions showed significant activation for the contrast of post-learning < pre-learning stages in the JRD task. **(d)** Brain regions showing the activation associated with *ED*_*ref-tar*_ in the JRD task. The warm/cold colors indicate positive/negative correlation. L/R indicates the left/right hemisphere. **(e)** Significant brain-behavior correlation. The *x-*axis represents the random effect coefficients (*η*) of the learning stage from the linear mixed-effects model (LMM), and the *y*-axis represents the *β*-values from the general linear model (GLM) analysis for the JRD trials and *ED*_*ref-tar*_. The coordinates indicate the centers of ROIs. The shaded area indicates the 95% confidence interval (CI). The detailed brain-behavior correlation for all brain regions involved in the JRD task are listed in Table S2 (Supplementary Data).

**Fig. 4.**
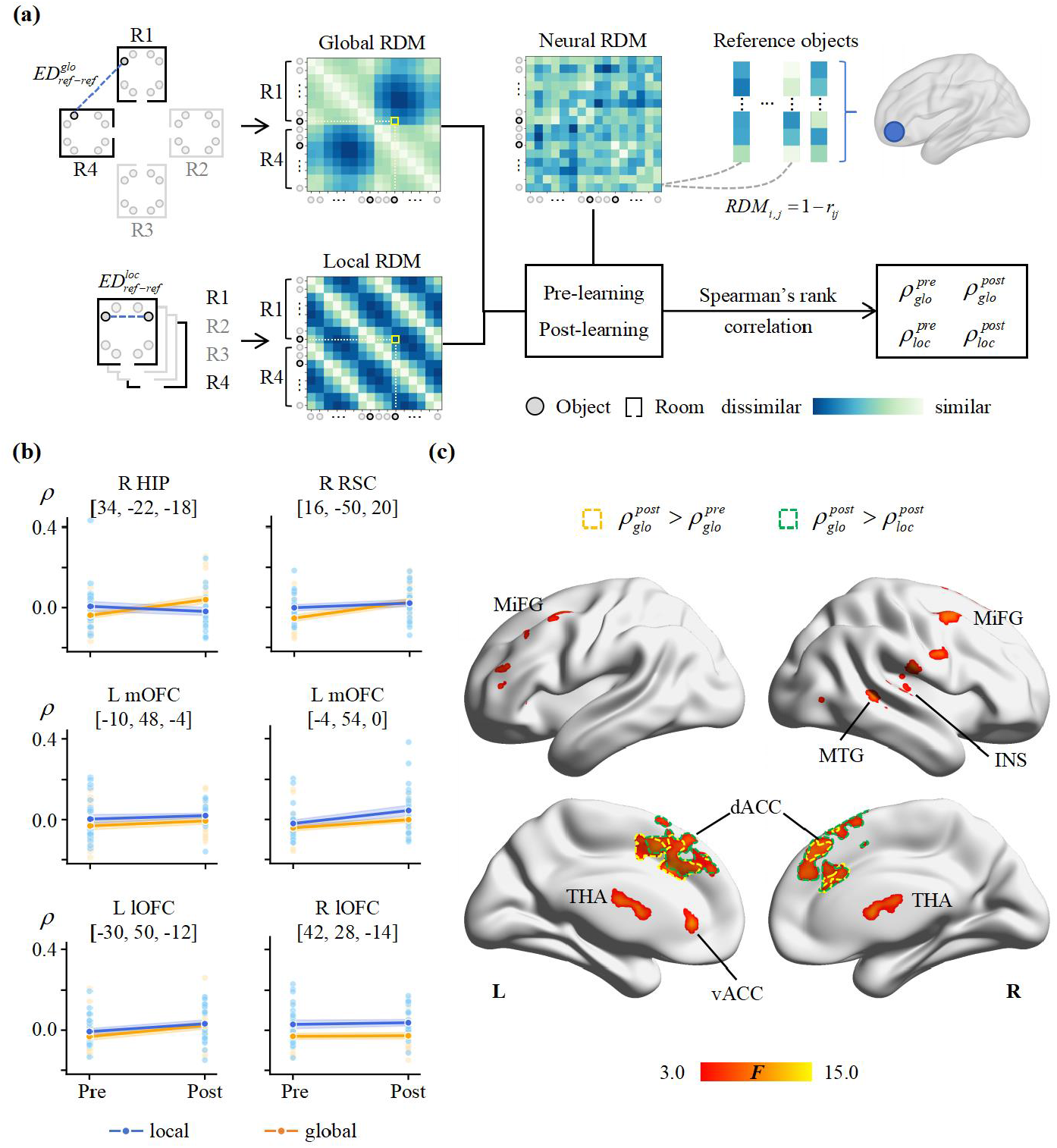
Representational similarity analysis (RSA). **(a)** Schematic of the RSA. Two theoretical representational dissimilarity matrices (RDMs), one for global and one for local levels, were built separately for both the learning stages (pre-learning and post-learning) to capture the spatial relationships of the reference objects. Considering that each room contained 8 objects and each learning stage used two rooms (16 objects total), we built the RDMs as 16 × 16 matrices. The left panel is an example of a pair of reference objects (highlighted in yellow in the matrices) in a single learning stage. Black (gray) rectangles indicate rooms that were (not) used in the learning stage. The global RDM and local RDM were built using the Euclidean distance between the reference objects within the global reference frame 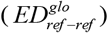 and the local reference frame 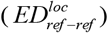, respectively, with larger distances indicating greater dissimilarity. For both the pre-learning and post-learning stages, the neural RDM was built by (1 - *r*_*ij*_), where *r*_*ij*_ represents Pearson’s correlation coefficient between the *β*-values for the *i*^th^ and *j*^th^ reference objects within a given brain region. Light (dark) colors in the matrices indicate high (low) similarity. Corresponding to the two hierarchical levels and two learning stages, four Spearman’s rank correlations were computed by estimating the correlation between the neural RDMs and each theoretical RDM. This yielded four correlation coefficients: 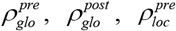, and 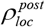 **(b)** Significant results of the ROI-based RSA. The coordinates represent the gravity centers of ROIs. Pre = pre-learning stage, Post = post-learning stage. **(c)** Brain regions showing significant differences in the correlation coefficients. Brain regions outlined by yellow and green dashed lines showed significant 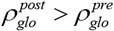 and 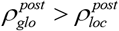 respectively. Abbreviations: HIP = hippocampus, RSC = retrosplenial cortex, l/mOFC = lateral/medial orbitofrontal cortex, d/vACC = dorsal/ventral anterior cingulate cortex, THA = thalamus, MiFG = middle frontal gyrus, INS = insula, MTG = middle temporal gyrus. L/R represents left/right hemisphere.

The neural RDM was calculated based on the fMRI data. For each trial, *β*-maps representing brain activity were computed using Nibetaseries toolbox (https://nibetaseries.readthedocs.io/en/v0.6.0/). The *β*-maps for each reference object were averaged across trials to create an overall *β*-map for that object. The neural RDM was then generated by calculating the dissimilarity (1 - *r*_*ij*_) between the *β*-values of different reference objects within each brain region, where *r*_*ij*_ is the Pearson’s correlation coefficient between the activation patterns for the *i*^th^ and *j*^th^ objects. This process was repeated for each brain region and for each of the two learning stages.

We conducted both the ROI-based and whole-brain searchlight RSA. In the ROI-based RSA, we created 4 mm-radius spherical ROIs centered at the peak coordinates of significant clusters which were identified in the GLM analysis. In addition, we used the bilateral M1 as the control ROIs. Neural RDMs were computed within these ROIs. For each subject at each learning stage, Spearman’s rank correlation (*ρ*) was calculated between each of the theoretical RDMs (local and global) and the neural RDM to assess the spatial representation in each ROI between the local and global levels (Fig. 4a). In the whole-brain searchlight RSA, we used a 4 mm-radius spherical searchlight and moved it across the brain with a stride of 1 voxel along each axis (*x, y, z*) to generate the whole-brain *ρ*-maps representing the correlation between the theoretical and neural RDMs for each subject in each learning stage. Fisher’s *ρ*-to-*Z*-transformation was applied to standardize the *ρ*-maps.

### 2.10 Statistics

For the group-level GLM analysis, we used a one-sample *t*-tests to assess the effects of the JRD trials and *ED*_*ref-tar*_ on brain activation. A paired-sample *t*-tests was conducted to evaluate the effects of learning on the task-related activation. Multiple comparisons were corrected using Gaussian Random Field (GRF) theory, with a voxel-wise threshold of *Z* > 3.1 and a cluster significance threshold of *p* < 0.05. For the RSA, each brain region yielded four Spearman’s rank correlations 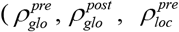, and 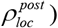, corresponding to the two hierarchical levels (global/local) and two learning stages (pre/post), which were computed between each of the neural RDMs and each of the theoretical RDMs. In the ROI-based RSA, a repeated measures ANOVA was performed to examine the effects of the hierarchical level and the learning stage on spatial representations within the ROIs, using a Python package, statsmodels (https://www.statsmodels.org/stable/index.html). For the whole-brain searchlight RSA, we performed a repeated measures ANOVA by using PALM (permutation analysis of linear models) to identify the brain regions where the spatial representation varied by the hierarchical level and the learning stage. Threshold-free cluster enhancement (TFCE) with 10,000 permutations and family-wise error (FWE) correction were used to control for multiple comparisons. The significant level was set at *p* < 0.01. A simple main effect analysis was conducted for significant ANOVA results. For the main effect of the learning stage on spatial representation, we tested whether the enhancement in spatial representation at each level after learning (i.e., 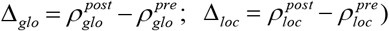 was significant. For the main effect of hierarchical levels on spatial representation, we tested the difference in spatial representation between the two levels in each learning stage (i.e.,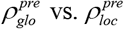 and 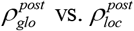 . In addition, we tested whether the post-learning improvement in spatial representation differed between the two hierarchical levels (i.e., Δ_*glo*_ vs. Δ_*loc*_ ). TFCE with 10,000 permutations and FWE correction were used to control for multiple comparisons in the main effect analyses, and the significant level was set at *p* < 0.01.

## 3 Results

### 3.1 Behavioral analysis using LMM

LMMs were used to analyze the influence of spatial information at different hierarchical levels on the subjects’ response in the two learning stages. Model comparison showed that LMM2 provided a better fit than LMM1, as indicated by the *AIC* values (*AIC*_LMM1_ = 5,737 > *AIC*_LMM2_ = 5,518), suggesting individual differences in the learning process. In the following, we reported the results obtained from LMM2.

Table 1 lists the main effects of different factors on RT, and their interactions with *Learning*. We found significant main effect of *Learning* (*F* = 39.61, *p* < .001), *ED*_*ref-tar*_ (*F* = 309.36, *p* < .001), 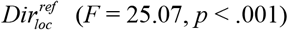, *Room* (*F* = 5.17, *p* = .023) and 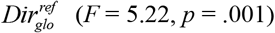 on RT. We also found significant interactions between *Learning* and *ED* (*F* = 6.03, *p* = .014) as well as between *Learning* and 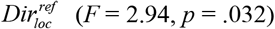 Specifically, the effect of *ED*_*ref-tar*_ on RT was stronger in the post-learning stage than in the pre-learning stage, suggesting that the subjects developed a better understanding of the spatial relationships between the objects after learning. Conversely, the effect of 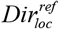 on RT was decreased after learning, indicating that the subjects may have formed internal representation of the environment (Table S1 in Supplementary Data). However, no significance was found in either the interaction between *Room* and *Learning* (*F* = 1.36, *p* = .244) or the interaction between 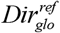 and *Learning* (*F* = 2.04, *p* = .106). This indicates that the influence of room alignment and global reference direction on RT was independent of the learning stages. For 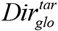 and 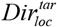, neither the main effects (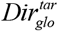 : *F* = 1.15, *p* = .329; 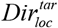: *F* = 2.58, *p* = .051) nor their interactions with *Learning* (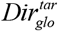 : *F* = 2.37, *p* = .069; 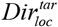 : *F* = 1.97, *p* = .116) reached the significance level.

**Table 1.**
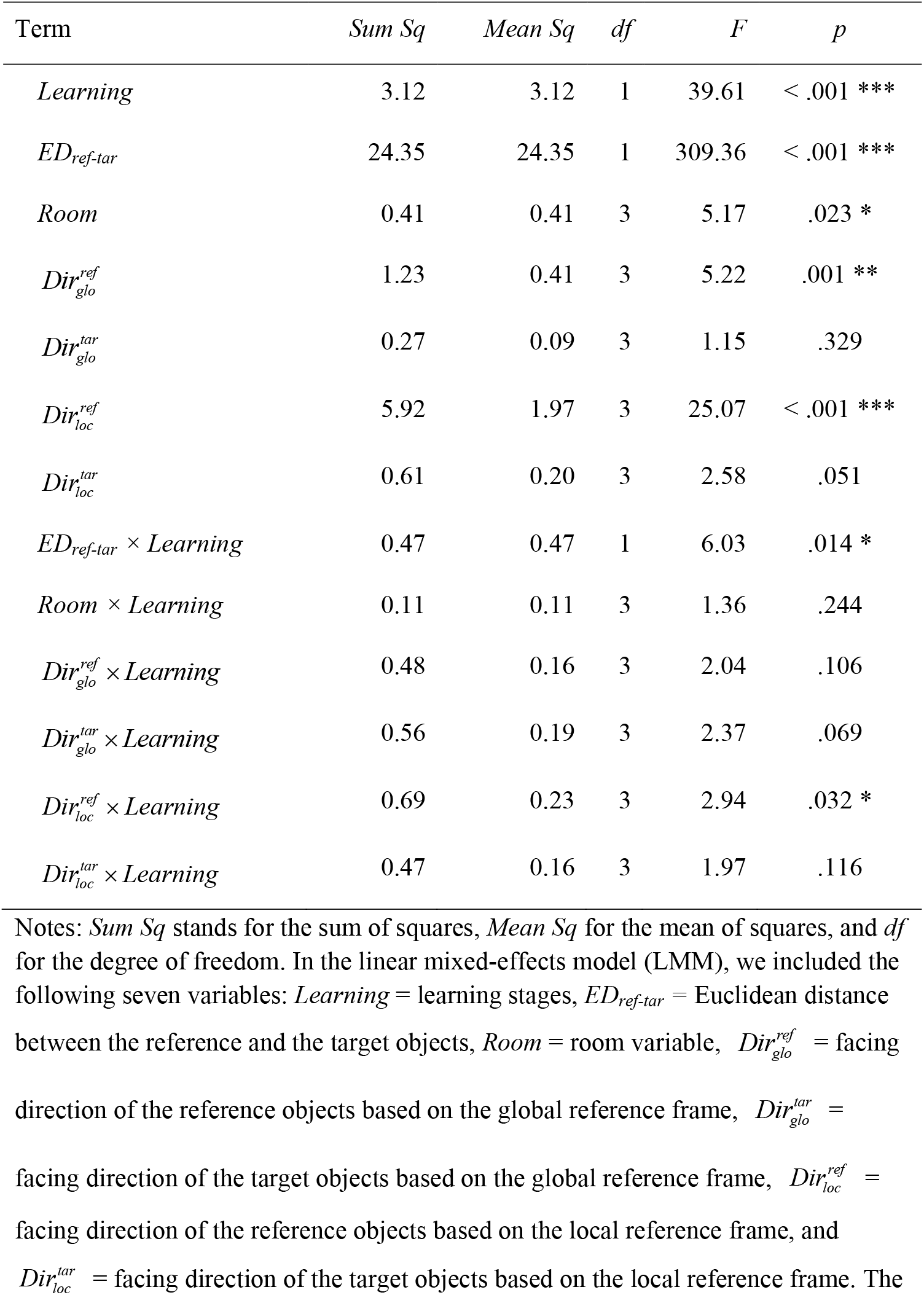
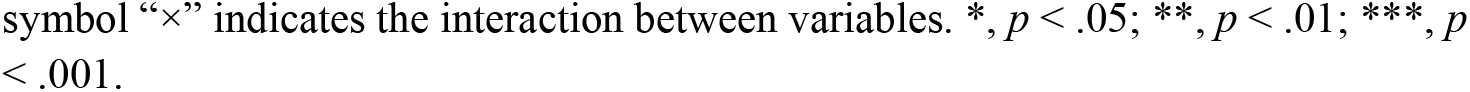
Fixed-effects ANOVA results obtained from the linear mixed-effects model (LMM).

| Term | *Sum Sq* | *Mean Sq* | *df* | *F* | *p* |
| --- | --- | --- | --- | --- | --- |
| *Learning* | 3.12 | 3.12 | 1 | 39.61 | < .001 *** |
| *ED<sub>ref-tar</sub>* | 24.35 | 24.35 | 1 | 309.36 | < .001 *** |
| *Room* | 0.41 | 0.41 | 3 | 5.17 | .023 * |
| *Dir<sub>glo</sub><sup>ref</sup>* | 1.23 | 0.41 | 3 | 5.22 | .001 ** |
| *Dir<sub>glo</sub><sup>tar</sup>* | 0.27 | 0.09 | 3 | 1.15 | .329 |
| *Dir<sub>loc</sub><sup>ref</sup>* | 5.92 | 1.97 | 3 | 25.07 | < .001 *** |
| *Dir<sub>loc</sub><sup>tar</sup>* | 0.61 | 0.20 | 3 | 2.58 | .051 |
| *ED<sub>ref-tar</sub> × Learning* | 0.47 | 0.47 | 1 | 6.03 | .014 * |
| *Room × Learning* | 0.11 | 0.11 | 3 | 1.36 | .244 |
| *Dir<sub>glo</sub><sup>ref</sup> × Learning* | 0.48 | 0.16 | 3 | 2.04 | .106 |
| *Dir<sub>glo</sub><sup>tar</sup> × Learning* | 0.56 | 0.19 | 3 | 2.37 | .069 |
| *Dir<sub>loc</sub><sup>ref</sup> × Learning* | 0.69 | 0.23 | 3 | 2.94 | .032 * |
| *Dir<sub>loc</sub><sup>tar</sup> × Learning* | 0.47 | 0.16 | 3 | 1.97 | .116 |

Notes: *Sum Sq* stands for the sum of squares, *Mean Sq* for the mean of squares, and *df* for the degree of freedom. In the linear mixed-effects model (LMM), we included the following seven variables: *Learning* = learning stages, *ED<sub>ref-tar</sub>* = Euclidean distance between the reference and the target objects, *Room* = room variable, *Dir<sub>glo</sub><sup>ref</sup>* = facing direction of the reference objects based on the global reference frame, *Dir<sub>glo</sub><sup>tar</sup>* = facing direction of the target objects based on the global reference frame, *Dir<sub>loc</sub><sup>ref</sup>* = facing direction of the reference objects based on the local reference frame, and *Dir<sub>loc</sub><sup>tar</sup>* = facing direction of the target objects based on the local reference frame. The
symbol “×” indicates the interaction between variables. \*, $p < .05$ ; \*\*, $p < .01$ ; \*\*\*, $p$ $< .001$ .

### 3.2 Brain activation during spatial memory

Fig. 3b shows seven clusters with significant activation and five clusters with significant deactivation in the JRD trials, which were identified from the ROI-based GLM analysis. The activated clusters were located in the bilateral RSC, EC, and lateral OFC (lOFC). The deactivated clusters were located in the bilateral medial OFC (mOFC), ventral lOFC, HIP, and EC. In the JRD trials, we found a cluster in the left RSC showing significant activation in post-learning > pre-learning (Fig. 3c). In addition, the activation in the left RSC was significantly positively correlated with *ED*_*ref-tar*_ (Fig. 3d). Conversely, the activation in the left mOFC, right HIP, and left lOFC was significantly negatively correlated with *ED*_*ref-tar*_ in the JRD trials. However, none of these correlations showed significant differences between the pre- and post-learning stages. The detailed information about these clusters is provided in Table S2 (Supplementary Data).

The whole-brain GLM analysis showed significant activation in the bilateral RSC and the left HIP for post-learning > pre-learning in the JRD task (Fig. S3, Supplementary Materials). We found that several regions, including the bilateral lateral occipital cortex (LOC), precuneus, supramarginal gyrus (SMG), left parahippocampal gyrus (PHG), and left middle temporal gyrus, showed significant activation for post-learning > pre-learning (Fig. S3, Supplementary Materials). The detailed information about these regions is listed in Table S3 (Supplementary Data).

Fig. 3e shows the brain-behavior correlations based on the random slopes of the learning stage from LMM2. Specifically, for the JRD trials, we found that the *β*-values of two right EC clusters (MNI peak: [28, -22, -28], *r* = -0.45, *p* = .032; MNI peak: [40, -6, -26], *r* = -0.50, *p* = .016) were significantly negatively correlated with the *η*-values. For the *ED*_*ref-tar*_, we found that the *β*-values of the right RSC (MNI peak: [14, -52, 20], *r* = 0.46, *p* = .029) and right HIP (MNI peak: [34, -22, -18], *r* = 0.66, *p* < .001) were significantly positively correlated with the *η*-values.

### 3.3 Spatial representation across learning stages

Fig. 4b shows the brain regions where spatial representations were significantly influenced by the hierarchical levels and/or the learning stages, as revealed by the ROI-based RSA. Specifically, we observed a significant interaction between the hierarchical level and the learning stage in the right HIP (MNI peak: [34, -22, -18], *F* = 6.34, *p* = .020). As learning progressed, the pattern similarity associated with the global RDM was increased, while that associated with the local RDM was decreased. This result suggests that the right HIP was more involved in representing the global spatial structure, contributing to the formation of a more integrated and comprehensive cognitive map of the VE.

In the right RSC (MNI peak: [16, -50, 20]), we also observed a significant interaction between the hierarchical level and the learning stage (*F* = 5.59, *p* = .027). Specifically, the enhancement in global representation after learning was greater than that in local representation (Fig. 4b). In addition, the main effect of learning stages on spatial representations was significant in the right RSC (*F* = 8.92, *p* = .007), with the pattern similarity increasing across both the local and global RDMs after learning. These results indicate the key role of the right RSC in integrating spatial information across different hierarchical levels.

Spatial representations in the OFC were influenced by both the hierarchical level and the learning stage. Specifically, in the left mOFC, we observed significant main effects of both the hierarchical level (MNI peak: [-10, 48, -4], *F* = 4.86, *p* = .038) and the learning stage (MNI peak: [-4, 54, 0], *F* = 9.35, *p* = .006). The pattern similarity of the local RDM was higher than that of the global RDM in both stages, and pattern similarity of both the RDMs were increased after learning (Fig. 4b). In the right lOFC (MNI peak: [42, 28, -14]), we observed a significant main effect of the hierarchical level (*F* = 21.30, *p* < .001), with the pattern similarity of the local RDM being stronger than that of the global RDM in both the pre- and post-learning stages. In the left lOFC (MNI peak: [-30, 50, -12]), we observed a significant main effect of the learning stage (*F* = 5.16, *p* = .033), with the pattern similarity of both the RDMs being increased after learning. No significant effects of the hierarchical level or the learning stage were found in the bilateral M1 (the control ROIs). The detailed information about these ROI-based RSA results is provided in Table S4 (Supplementary Data).

The whole-brain searchlight RSA revealed additional brain regions showing significant changes in spatial representations between the pre- and post-learning stages. Using ANOVA, we identified 13 brain clusters with different pattern similarity for the local and global RDMs across the two learning stages (Fig. 4c). These clusters were located in the bilateral ACC, bilateral thalami, bilateral lateral prefrontal cortex (lPFC), left OFC, right middle temporal gyrus (MTG), right lateral occipital cortex (LOC), right post-central gyrus (PoCG), and right pre-central gyrus (PrCG) (Table S5 in Supplementary Data). In the post-learning stage, these regions showed higher pattern similarity for the global RDM than in the pre-learning stage and higher similarity for the global RDM than the local RDM (Fig. S4 in Supplementary Materials). However, only the bilateral ACC showed these differences to be statistically significant (Fig. 4c). Both the bilateral ACC and lPFC showed reduced pattern similarity for the local RDM after learning. These regions also showed lower pattern similarity for the global RDM than the local RDM in the pre-learning stage (Fig. S4 in Supplementary Materials). However, neither of these results reached the significant level.

When comparing the two learning stages, we observed significant differences in the local and global spatial representations in four clusters. These clusters were located in the bilateral ACC, left inferior frontal gyrus (IFG), and right superior frontal gyrus (SFG), with the corresponding details provided in Fig. S5 (Supplementary Materials). In these clusters, the post-learning enhancement in pattern similarity of the global RDM was greater than that of the local RDM ( Δ_*glo*_ > Δ_*loc*_ ). The detailed information about these searchlight RSA results is provided in Table S5 (Supplementary Data).

## 4. Discussion

The current study examined how cognitive maps develop in a multi-scale environment through learning. We used a virtual environment (VE) that included spatial information at three hierarchical levels: local, categorical, and global (Fig. 2), allowing the subjects to learn the spatial relationships between objects. Their spatial memory was measured during both pre-learning and post-learning stages using a JRD task (Fig. 1). By analyzing the behavioral data, we found that as learning progressed, the subjects gradually shifted from relying on explicit directional cues to processing implicit spatial distance (Fig. 3a), indicating a refinement of cognitive map. By analyzing the fMRI data, we revealed that the HIP, EC, RSC, and OFC were engaged in encoding spatial information during the JRD task (Fig. 3). These regions played distinct roles in representing spatial information at the local and global levels in the pre-and post-learning stages (Fig. 4). Specifically, learning enhanced global representations in the HIP, strengthened local representations in the OFC, and increased representations at both hierarchical levels in the RSC (Fig. 4b). These findings indicate the different roles of these key brain regions in supporting the dynamic construction of cognitive maps. By uncovering how hierarchical spatial representations emerge and re-organize with learning, the current study advances our understanding of the neural basis of flexible navigation in complex environments.

### 4.1 Refinement on cognitive map through learning

The LMM analysis revealed significant improvements in behavioral performance from the pre-learning to the post-learning stage (Table 1). After learning, the subjects showed a better understanding of the spatial relationships between objects, as indicated by the increased influence of *ED*_*ref-tar*_ on RT in the post-learning stage than the pre-learning stage (Fig. 3a). In addition, the influence of categorical information (i.e., the *Room*) on RT was decreased after learning (Table S1, Supplementary Data), indicating a reduced reliance on room-level distinctions as the subjects became more familiar with the environment. These results are consistent with previous findings ^21,22^, which showed that individuals tend to shift from using explicit categorical information to using more implicit spatial representations during spatial learning.

How did local and global representations evolve throughout the learning process? From the LMM, we found a significant influence of the local directions 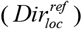 and global directions 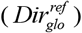 of the reference objects on RT in both the pre- and post-learning stages (Table 1). This indicates that both local and global representations co-existed in both the learning stages. However, the influence of these directional cues on the representation was reduced in the post-learning than the pre-learning stages (Fig. 3a), suggesting a decreased reliance on the explicit directional cues as learning progressed. These findings align with previous studies ^16,20^, which showed that both local and global representations co-exist in multi-scale environments and are recruited flexibly according to task demands.

### 4.2 Neural mechanisms in the JRD task facilitating spatial learning

The GLM analysis revealed that the bilateral HIP, EC, RSC, and OFC were engaged in the JRD task (Fig. 3b). These regions are involved in various cognitive functions, such as learning, memory, spatial navigation, and cognitive maps formation ^1,25,26^. In the current study, we found that the left RSC showed significantly greater activation in the post-learning stage than in the pre-learning stage (Fig. 3c). This result aligns with a previous research ^35^, which showed the critical role of the RSC in building cognitive maps of novel environments. This result indicates that the RSC may be involved in processing spatial relationships across different scales of the environment ^33,43^. In addition, we found that the individuals with greater learning progress showed weaker activation in the right EC (Fig. 3e). The EC is proposed encoding general, non-localized spatial information across different environmental scales ^2,6^. This negative correlation between EC activation and learning progress suggests that spatial representations may be transferred from the EC to other brain regions with the progress of learning, leading to a reduced reliance on the EC ^44,45^.

We also observed that activation in the left mOFC, right lOFC, right HIP, and right RSC was significantly correlated with *ED*_*ref-tar*_ in the JRD task (Fig. 3d). This result is consistent with previous studies ^24,46,47^, which showed that the HIP and RSC are involved in encoding distances in both physical and abstract spaces. The mOFC and HIP are suggested to contribute to the construction of cognitive maps that support flexible decision-making and adaptive behavior ^2,5,15^. In addition, we found the positive correlation between activation in the RSC and HIP with the progress of individuals’ learning (Fig. 3e). This result suggests that the RSA and HIP may support the formation of individual-specific spatial representations ^25,35^.

### 4.3 Adaptive shifts in spatial representations during learning

The RSA results revealed that the brain regions involved in the JRD task played distinct roles in the formation of a cognitive map in a multi-scale environment (Fig. 4). Specifically, in the right HIP, we observed a shift from a local representation in the pre-learning stage to a global representation in the post-learning stage (Fig. 4b). This observation aligns with previous studies ^35,48,49^, which showed that the HIP is crucial for integrating local information into a global layout as learning progresses. In the RSC, spatial representations of both local and global levels were increased from pre-learning to post-learning (Fig. 4b), indicating the RSC’s role in integrating spatial information across spatial scales. This aligns with previous studies ^25,32,33^, which implicated the roles of RSC in representing location and facing direction in multi-scale environments. Similarly, both local and global representations were increased in the left OFC in the post-learning stage (Fig. 4b), suggesting that it coordinates spatial information across different scales, and supporting adaptive decision-making and goal-directed behavior ^34,50,51^. In contrast, the right lOFC consistently showed strong local representation in both the pre and post-learning stages (Fig. 4b), indicating its role in processing specific location and direction information for decision-making ^30,31,50^.

Using whole-brain searchlight RSA, we found additional brain regions showing changes in spatial representations across the two learning stages (Fig. 4c). Specifically, the bilateral ACC showed an increase in global representation after sufficient learning (Fig. 4c). The enhancement of global representations in the ACC was more pronounced than that of local representations (Fig. S5 in Supplementary Materials), resulting in a higher global representation than local representation after learning (Fig. 4c). Previous studies indicated that the ACC is functionally connected to the OFC and HIP in spatial memory, especially in tasks relating to exploration in novel environments ^52–54^. Our results suggest that the ACC may be engaged in integrating spatial information to form global representations, thereby facilitating efficient and flexible spatial processing. In addition, other brain regions, including the bilateral lPFC, bilateral thalami, right insula, left ventral ACC, and right MTG, showed varying spatial representations across the learning stages (Fig. 4c). In these regions, both global and local representations were increased after sufficient learning, with global representations remaining consistently stronger than local representations across both learning stages (Fig. S4 in Supplementary Materials). Though these regions did not show statistical significant main effects of spatial hierarchical levels or learning stages, they have been implicated in various learning processes and are associated with the HIP in explicit memory ^55–57^. Our results suggest that forming cognitive maps in multi-scale environments recruits multiple brain regions to integrate information across spatial levels.

### 4.4 Disentangling Global and Categorical Representations

We observed an increase in global representations after sufficient learning, which could arise from two potential mechanisms. First, the subjects may have acquired precise spatial relationships between the rooms and objects. This hypothesis is supported by the LMM results, which showed that the distance information became more influential in behavioral performance after learning (Fig. 3a). Second, the increase in global representation could be resulted from the improved differentiation of rooms (i.e., categorical processing), rather than the acquisition of precise spatial distances. In this case, the subjects may have enhanced their categorical-level spatial representation, which does not require the precise distance relationships inherent to the global representation.

Categorical representations are essential for distinguishing boundaries and characteristics between areas, enabling transitions across different areas in multi-scale environments ^17^. These representations lie between the local and global representations in the hierarchical representation framework (Fig. 2). Unlike the global representation, which conveys precise distance relationships between objects, the local and categorical representations are less precise. Local representations assimilate rooms by treating the distance between them as zero, and categorical representations differentiate rooms by treating the inter-room distance as infinite. Our LMM analysis indicated that the categorical information had a reduced influence on the behavioral performance after learning (Fig. 3a). This result suggested that the increased global representation observed in the RSA may be driven by the acquisition of precise distance information rather than the acquisition of categorical information. However, the high similarity between the global and categorical RDMs (Fig. S4 in Supplementary Materials) makes it challenging to distinguish brain representations at these two hierarchical levels. Therefore, further studies are needed to understand how spatial representations evolve across different hierarchical levels through learning.

### 4.5 Limitations

The current study has several limitations. First, we used a virtual reality (VR) environment to simulate spatial learning and navigation. Although VR offers a valuable tool for investigating spatial cognition ^58,59^, it may not fully capture the complexities and sensory richness of real-world environments. Sensory cues such as proprioceptive feedback and natural visual information, which are limited in VR settings, can influence the subjects’ spatial encoding strategies ^60^ and their reliance on the local and global spatial information. Second, our study employed a cross-sectional design, measuring subjects at two time points (pre- and post-learning). Although this design allows us to observe changes in brain activation and behavioral performance, it does not provide a direct measure of the process of learning over time. Longitudinal studies tracking the trajectory of individual learning would provide deeper insights into the dynamic nature of spatial learning and cognitive map development. Third, although we accounted for individual differences in the LMM analysis, variations in cognitive strategies and previous experiences with spatial tasks could influence how individuals process spatial information. Future studies could explore which factors, such as prior spatial experience, cognitive abilities, and personality traits, influence the progress of learning and the representation of spatial information. Last, our sample consisted of 23 adult healthy university students, their age range and educational levels may limit the generalizability of the findings ^61,62^. A larger and more diverse sample would help to test the observed effects.

In conclusion, the current study provides new insights into the neural mechanisms underlying spatial learning and the formation of cognitive maps in multi-scale environments. Using a combination of behavioral data and neuroimaging techniques, we showed how spatial representations evolve across two different learning stages. Specifically, we found that as learning progressed, individuals increasingly relied on global representations of the environment, with a shift away from local, directional cues. Brain regions including the hippocampus, retrosplenial cortex, and orbitofrontal cortex played distinct roles in encoding spatial information at multiple hierarchical levels, supporting the development of flexible and adaptive cognitive maps. These results revealed the dynamic nature of spatial learning and the distinct roles of the hippocampus, retrosplenial cortex, and orbitofrontal cortex in integrating local and global spatial information over time. The findings advance our understanding of the neural mechanisms of spatial learning in complex environments.

## Acknowledgements

This work was supported by funding from the Key Technologies R&D Program of Guangdong Province (Grant number: 2023B0303020002), National Natural Science Foundation of China (Grant numbers: 32371101 and 82171914), striving for the first-class, improving weak links and highlighting features (SIH) key discipline for psychology in South China Normal University, and National Key Research and Development Program of China (Grant number: 2018YFC1705006). We thank the Center for Magnetic Resonance Research (CMRR) at the University of Minnesota for providing the SMS-EPI sequence used in the current study.

## Notes

### Competing Interest Statement

The authors have declared no competing interest.

